# GeTallele: a method for integrative analysis and visualization of DNA and RNA allele frequencies

**DOI:** 10.1101/491209

**Authors:** Piotr Słowiński, Muzi Li, Paula Restrepo, Nawaf Alomran, Liam F. Spurr, Christian Miller, Krasimira Tsaneva-Atanasova, Anelia Horvath

## Abstract

**Background:** Asymmetric allele expression typically indicates functional and/or structural features associated with the underlying genetic variants. When integrated, RNA and DNA allele frequencies can reveal patterns characteristic of a wide-range of biological traits, including ploidy changes, genome admixture, allele-specific expression and gene-dosage transcriptional response.

**Results:** To assess RNA and DNA allele frequencies from matched sequencing datasets, we introduce a method for generating model distributions of variant allele frequencies (VAF) with a given variant read probability. In contrast to other methods, based on whole sequences or single SNV, proposed methodology uses continuous multi-SNV genomic regions. The methodology is implemented in a GeTallele toolbox that provides a suite of functions for integrative analysis, statistical assessment and visualization of **Ge**nome and **T**ranscriptome **allele** frequencies. Using model VAF probabilities, GeTallele allows estimation and comparison of variant read probabilities (VAF distributions) in a sequencing dataset. We demonstrate this functionality across cancer DNA and RNA sequencing datasets.

**Conclusion:** Based on our evaluation, variant read probabilities can serve as a dependable indicator to assess gene and chromosomal allele asymmetries and to aid calls of genomic events in matched sequencing RNA and DNA datasets.

## 1 Introduction

RNA and DNA carry and present genetic variation in related yet distinct manners; the differences being informative of functional and structural traits. In diploid organisms, an important measure of genetic variation is the allele frequency, which can be measured from both genome (DNA) and transcriptome (RNA) sequencing data (encoded and expressed allele frequency, respectively). Differential DNA-RNA allele frequencies are associated with a variety of biological processes, such as genome admixture or allele-specific transcriptional regulation (Ferreira, et al., 2016; Ha, et al., 2012; Han, et al., 2015; Movassagh, et al., 2016; Shah, et al., 2012).

Most of the RNA-DNA allele comparisons from sequencing have been approached at nucleotide level, where it has proven to be highly informative for determining the alleles’ functionality (Ferreira, et al., 2016; Ha, et al., 2012; Han, et al., 2015; Macaulay, et al., 2016; Morin, et al., 2013; Movassagh, et al., 2016; Reuter, et al., 2016; Shah, et al., 2012; Shi, et al., 2016; Shlien, et al., 2016; The, et al., 2012; Yang, et al., 2016). Comparatively, integration of allele signals at the molecular level, as derived from linear DNA and RNA carriers, is less explored due to challenges presented by short sequencing length. The different molecular nature of RNA and DNA also leads to limited compatibility of the sequencing output.

Herein, we address some of the above challenges by employing a mathematical model to assess differences between RNA- and DNA-variant allele frequencies (VAF) at single nucleotide variant (SNV) positions in the genome. To do that, we introduce GeTallele: a toolbox that provides a suit of functions for integrative analysis and visualization of **Ge**nome (VAF_DNA_) and **T**ranscriptome (VAF_RNA_) **allele** frequencies along genes and chromosomes. Using VAF, GeTallele infers possible copy number alterations (CNAs), and, at positions of interest, mathematically and statistically compares VAF_RNA_ and VAF_DNA_ distributions. Furthermore, GeTallele supports visualization of the allele distribution at a desired resolution—from nucleotide to genome. In contrast to other CNA-modelling methods based on statistical models, GeTallele is based on a mechanistic model of VAF distributions.

## 2 Results

We demonstrate GeTallele’s functionality using sequencing DNA and RNA datasets from paired normal and tumour tissue obtained from 72 female patients with breast invasive carcinoma (BRCA) from The Cancer Genome Atlas (TCGA). Each dataset contains four matched sequencing sets: normal exome (Nex), normal transcriptome (Ntr), tumour exome (Tex), and tumour transcriptome (Ttr). The raw sequencing data were processed as previously described (Movassagh, et al., 2016) to generate the inputs for GetAllele.

The overall workflow of GetAllele is shown on Figure 1. As an input, GeTallele requires the absolute number of sequencing reads bearing the variant and reference nucleotide in each single-nucleotide variant (SNV) position across the four matched datasets. For each sample, to select SNV positions for analysis, we start with the high quality heterozygous SNV calls on the normal exome (Li, et al., 2009). In each of these positions we estimate the counts of the variant and reference reads (n_VAR_ and n_REF_, respectively) across the 4 matching datasets, and retain for further analyses only positions covered by a minimum total (variant and reference) required unique sequencing reads across. This threshold is flexible and is required to ensure that only sufficiently covered positions will be analysed; it is set to 3 in the herein presented results.

**Fig. 1.**
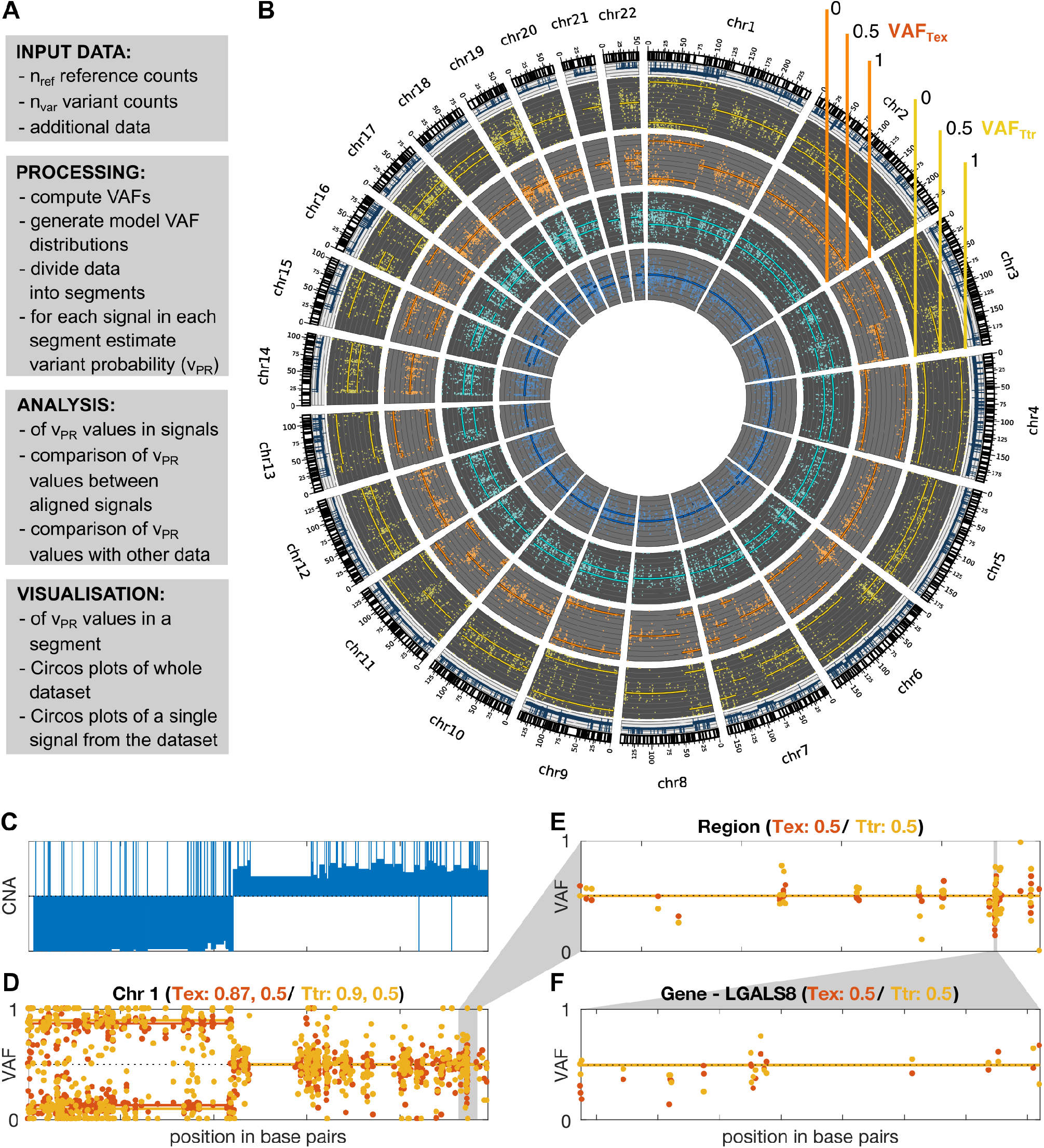
GeTallele and visualisation of VAF data. **A** Toolbox description. **B** Visualisation of the whole dataset on the level of genome using Circos plot (blue – normal exome, cyan – normal transcriptome, orange - tumour exome, yellow - tumour transcriptome). **C - F** show in details VAF_TEX_ and VAF_TTR_ values of chromosome 1; C - F Visualization of the VAF values with fitted variant probability (_VPR_ – see Section 3.1 and Figure 2) values at the level of chromosome (**D**), custom genome region (**E**) and gene (**F**), for the chromosome level shown also are CNA values (**C**). Panel **D** shows that there are two segments with different VAF distributions. Panel C shows that change in the CNA is concurrent with the change in the VAF distributions. Tex - tumour exome (orange); Ttr - tumour transcriptome (yellow).

From each of the 4 matched datasets GetAllele estimates VAF based on nVAR and nREF covering the positions of interest: VAF = n_VAR_/(n_VAR+nREF_). An example of genome-wide VAF values estimated from Tex, and their corresponding distribution of VAF values is shown in Figure 2.

**Fig. 2.**
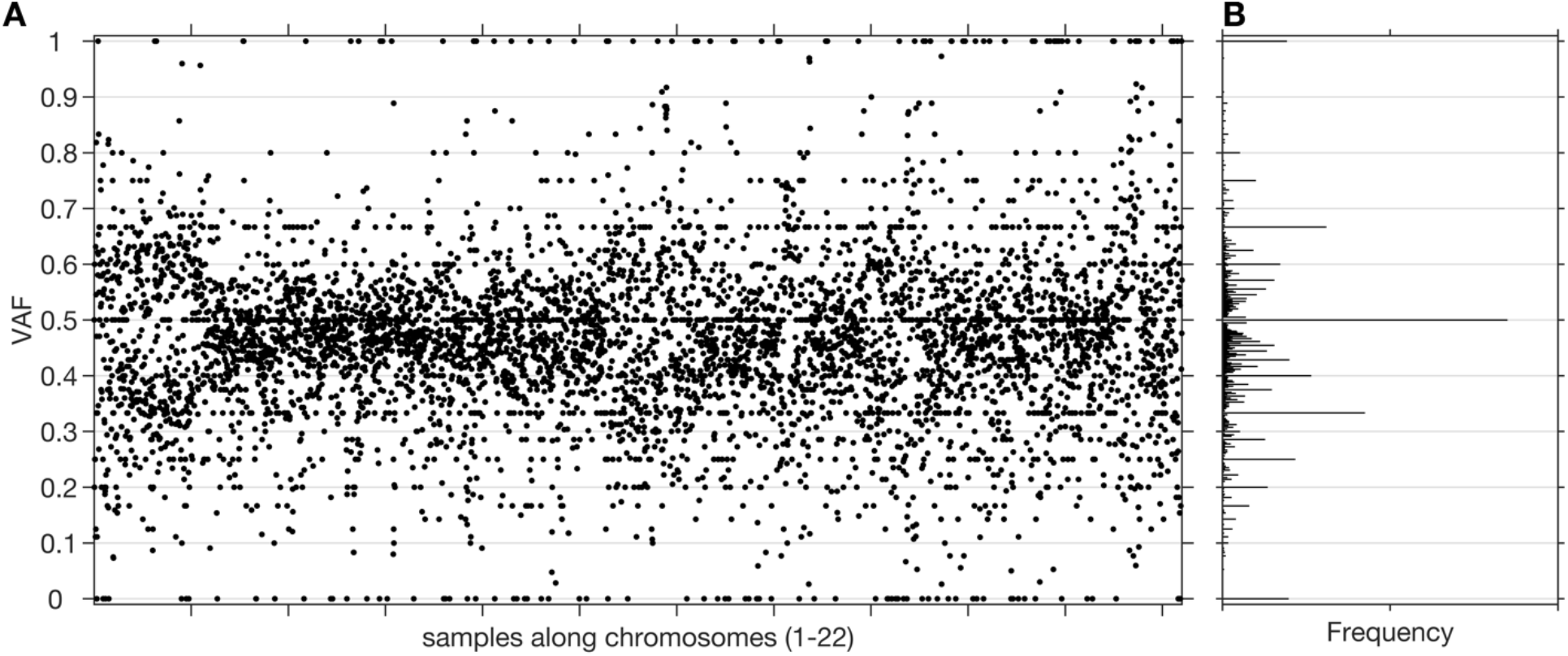
Sample and distribution of variant allele frequencies (VAF) values. **A.** All the VAF values of a tumour exome sequencing signal (chromosomes 1-22) from one of the datasets. **B.** Histogram of the VAF values from panel A. Centres of bins of the histogram are located at elements of a Farey sequence.

Analysis of the VAF_RNA_ and VAF_DNA_ in the GeTallele is based on comparing probability of observing a given VAF value at various positions of interest. Estimation of the variant allele probability (V_PR_) is implemented using a data-driven mathematical model of distribution of the VAF values and is the core functionality of the GeTallele.

### 2.1 Estimation of variant probability V_pr_

#### 2.1.1 Variant probability

GetAllele aggregates frequency counts of a VAF sample into variant probability, V_PR_ – probability of observing a variant allele. The V_PR_ is a single informative value indicative of copy number variation. The V_PR_ is a parameter that describes the genomic event that through the sequencing process was transformed into a specific distribution of VAF values found in the signal. For example, in VAF_DNA_ from a diploid genome, variant probability V_PR_=0.5 (meaning that both alleles are equally probable) corresponds to a true allelic ratio of 1:1 for heterozygote sites. For heterozygote sites in the DNA from diploid monoclonal sample, the corresponding tumour VAF_DNA_ is expected to have the following interpretations: V_PR_=1 or V_PR_=0 corresponds to a monoallelic status resulting from a deletion, and V_PR_=0.8 (or 0.2), 0.75 (or 0.25), 0.67 (or 0.33) correspond to alleles-pecific tetra-, tri-, and duplication of the variant-bearing allele, respectively.

The V_PR_ of the VAF_RNA_ is interpreted as follows. In positions corresponding to DNA heterozygote sites, alleles not preferentially targeted by regulatory traits are expected to have expression rates with variant probability V_PR_=0.5, which (by default) scale with the DNA allele distribution. Differences between VAF_DNA_ and VAF_RNA_ values are observed in special cases of transcriptional regulation where one of the alleles is preferentially transcribed over the other. In the absence of allele-preferential transcription, VAF_DNA_ and VAF_RNA_ are anticipated to have similar V_PR_ across both diploid (normal) and aneuploid (affected by CNAs) genomic regions. Consequently, VAF_DNA_ and VAF_RNA_ are expected to synchronously switch between allelic patterns along the chromosomes, with the switches indicating break points of DNA deletions or amplifications.

Since we observed that DNA and RNA signals have different distributions of total reads and also that the distributions of total reads vary between participants, the model VAF distributions are generated individually for each sequencing signal and each participant.

To estimate V_PR_ in the signals, GeTallele first generates model VAF distributions and then uses the earth mover’s distance (EMD) (Kantorovich and Rubinstein, 1958; Levina and Bickel, 2001) to fit them to the data. To generate a model VAF distribution with a given variant probability, V_PR_, GeTallele, bootstraps 10000 values of the total reads (sum of the variant and reference reads; n_VAR_ + n_REF_) from the analysed signal in the dataset. It then uses binomial pseudorandom number generator to get number of successes for given number of total reads and a given value of V_PR_ (implemented in the Matlab function `binornd`). The V_PR_ is the probability of success and generated number of successes is interpreted as an n_VAR_. Since the model V_PR_ can take any value, it can correspond to a single genomic event as well as any combination of genomic events in any mixture of normal and tumour populations (See section 2.1.3).

The analysis presented in the paper uses 51 model VAF distributions with V_PR_ values that vary from 0.5 to 1 with step (increment of) 0.01. The model VAF distributions are parametrized using only V_PR_ ≥ 0.5, however, to generate them we use V_PR_ and its symmetric counterpart 1-V_PR_. The process of generating model VAF distributions and examples of model and real VAF distributions with different values of V_PR_ are illustrated in Fig. 3.

**Fig. 3.**
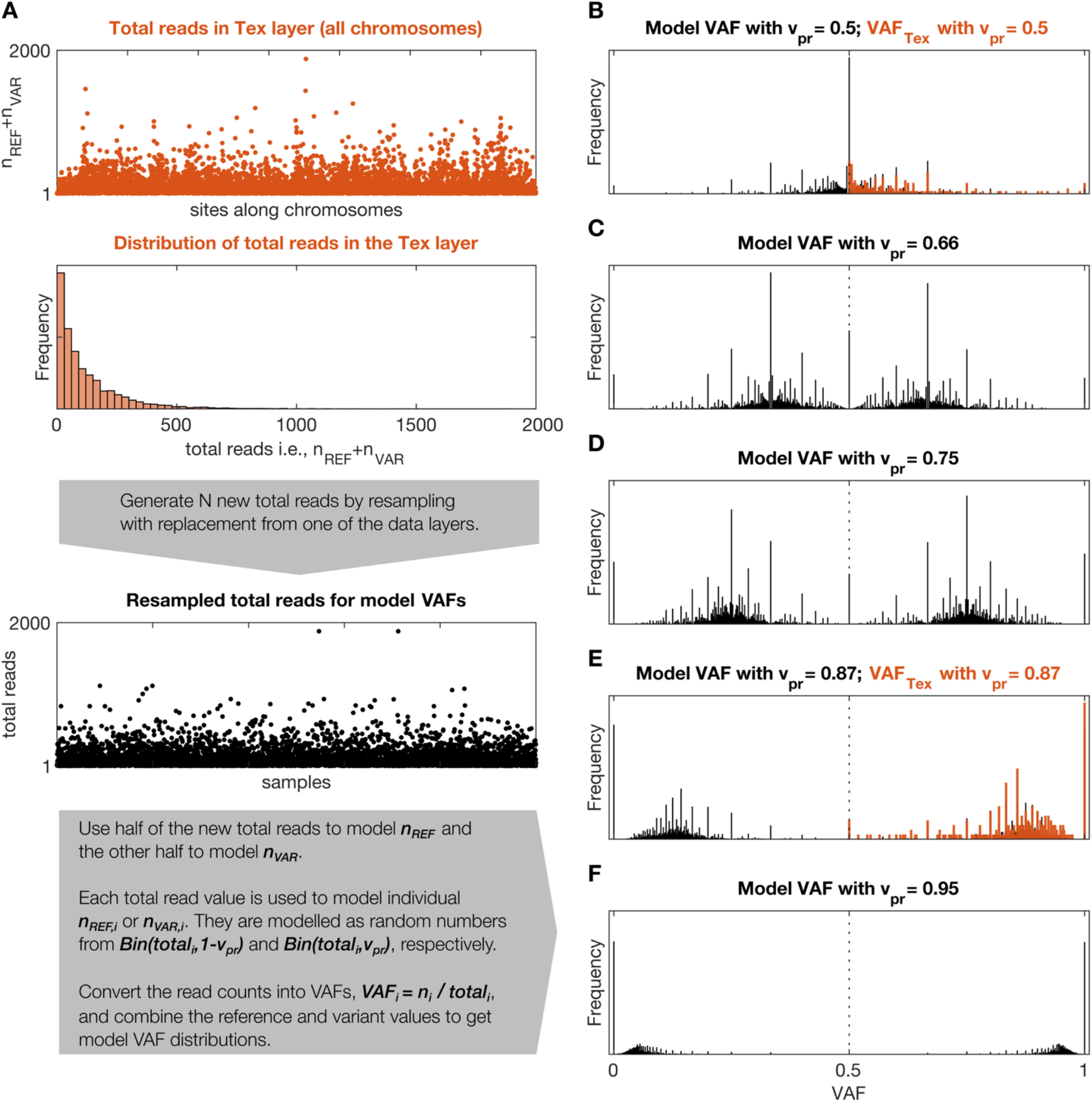
Model and real VAF distributions. **A** Description of the model used to generate model VAF distributions, *Bin(n,p)* stands for binomial distribution with parameters *n* (number of trials) and *p* (probability of success). **B** - **F** Model VAF distributions for different values of V_PR_. Panels **B** and **F** show additionally distributions of VAF_TEX_ for the two windows shown in Figure 1D.

#### 2.1.2 V_PR_estimation

Earth mover’s distance (EMD) is a metric for quantifying differences between probability distributions (Kantorovich and Rubinstein, 1958; Levina and Bickel, 2001) and in the case of univariate distributions it can be computed as:

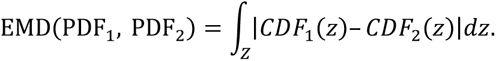

Here, PDF1 and PDF2 are two probability density functions and CDF_1_ and CDF_2_ are their respective cumulative distribution functions. Z is the support of the PDFs (i.e. set of all the possible values of the random variables described by them). Because VAFs are defined as simple fractions with values between 0 and 1, their support is given by a Farey sequence (Hardy, et al., 2008) of order n; n is the highest denominator in the sequence. For example, Farey sequence of order 2 is 0, 1/2, 1 and Farey sequence of order 3 is 0, 1/3, 1/2, 2/3, 1. GeTallele uses a Farey sequence of order 1000 as the support Z for all computations involving EMD.

To estimate V_PR_, GeTallele computes EMD between the distribution of the VAF values of each signal in the window and the 51 model VAF distributions (i.e. observed vs modelled VAF), the estimate is given by the V_PR_ of the model VAF distribution that is closest to the VAF distribution in the window. Examples of VAF distributions with fitted model VAF distributions are shown in Figure 3A and D. The dependence of the confidence intervals of the estimation on the number of VAF values in a window is illustrated in Fig. 4, which clearly demonstrates that the larger the number of VAFs in the chosen window the higher the accuracy of the estimate.

**Fig. 4.**
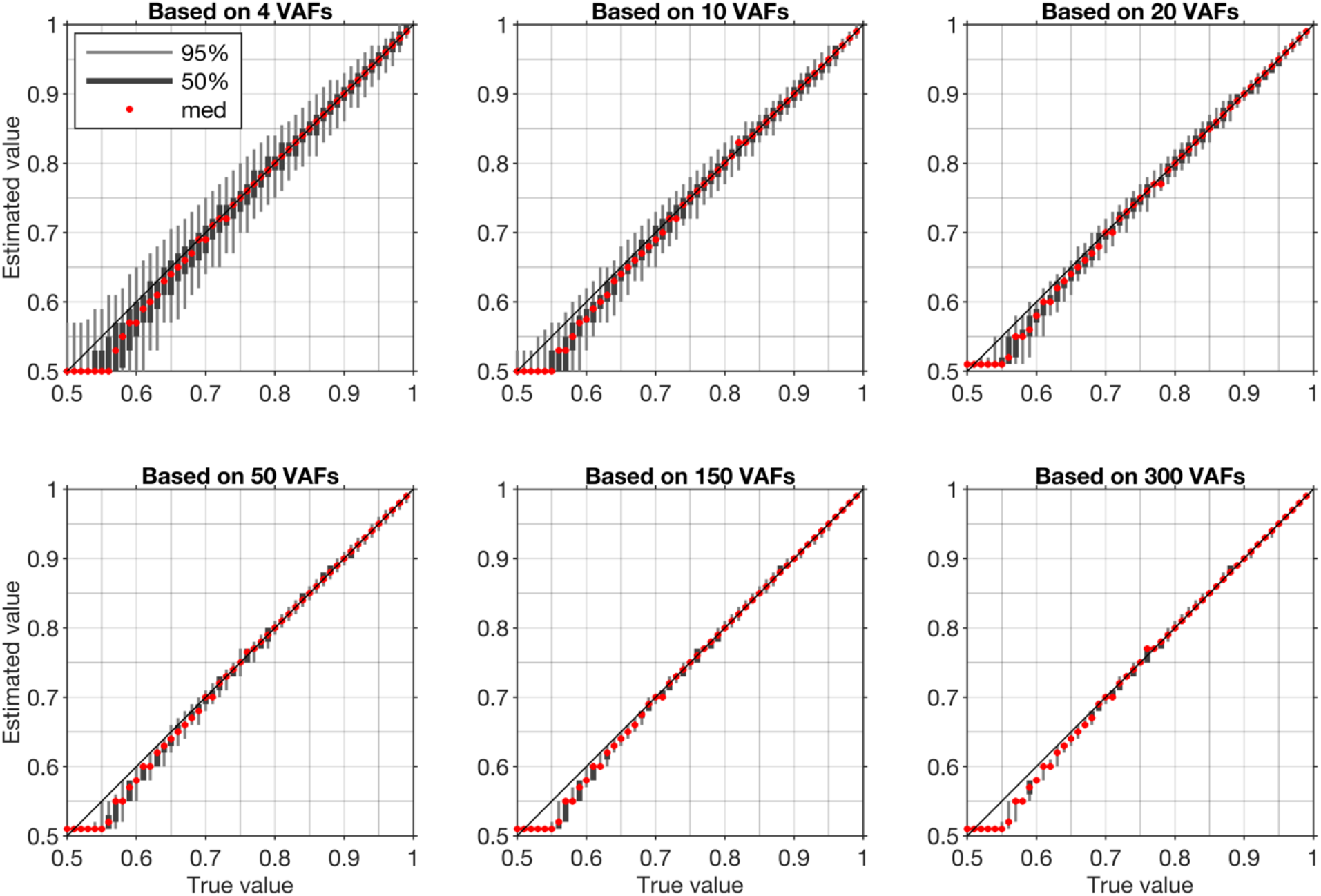
Confidence intervals for artificial samples with different numbers of VAFs. Each confidence interval is based on estimation of V_PR_ in 1000 randomly generated samples with a fixed V_PR_ (True value). Light grey bar is 95% confidence interval (950 samples lay within this interval), dark grey bar is 50% confidence interval (500 samples lay within this interval), red dot is median value.

#### 2.1.3 V_PR_ values in mixtures of normal and tumour populations

Since the V_PR_ can take any value between 0.5 and 1 it can correspond to a single genomic event as well as any combination of genomic events in any mixture of normal and tumour populations. A mixture V_PR_ value that corresponds to a combination of genomic events can be computed using the following expression:

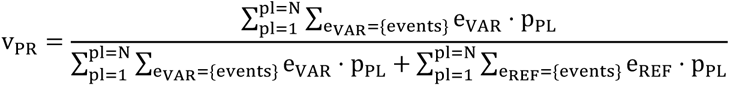

Where e_VAR_ and e_REF_ are the multiplicities of variant and reference alleles and p_PL_ is a proportion of one of the populations. For heterozygote sites e_VAR_=1 and e_REF_=1, for deletions e_VAR_=0 or e_REF_=0, for du-, tri- and tetraplications e_VAR_ or e_REF_ can be equal to 2, 3 or 4, respectively. The sum of proportions pPL over the populations is equal 1. For example, for a mixture of 1 normal (N, p_N_=0.44) and 2 tumour populations (T1, p_T1_=0.39 and T2, p_T2_=0.17), T1 with deletion and T2 with deletion the mixture V_PR_ value can be computed as follows:

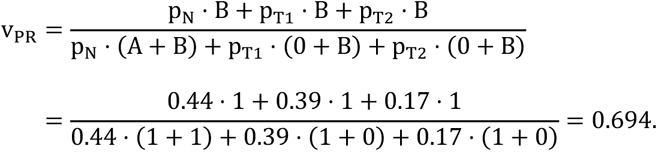

By comparing the V_PR_ values estimated from data with possible mixture V_PR_ values we propose to estimate sample purity and its clonal composition. To this end, we first generate a full set of proportions of all the population in the mixture with step (increment of) 0.01 and compute all the possible V_PR_ values that each of the mixtures could produce. For step 0.01: two populations (1 tumour) give 99 proportions, three populations (2 tumours) give 4851 proportions, four populations (3 tumours) give 156849 proportions. The matrices with mixture V_PR_ values for each proportion, vary from 2×2, for two populations with deletions, to 35×35 for four populations with all events up to tetraplications. Then, we run an exhaustive approximate search over all the matrices with mixture V_PR_ values over all the proportions. The search is approximate because the estimated V_PR_ values have limited accuracy and because we consider only discrete values of proportions. In the analysis we define a match between estimated and mixture V_PR_ values if they differ <0.009. The search returns a large number of admissible mixtures that could produce the estimated V_PR_ values. This process is illustrated in Fig. 5.

**Fig. 5.**
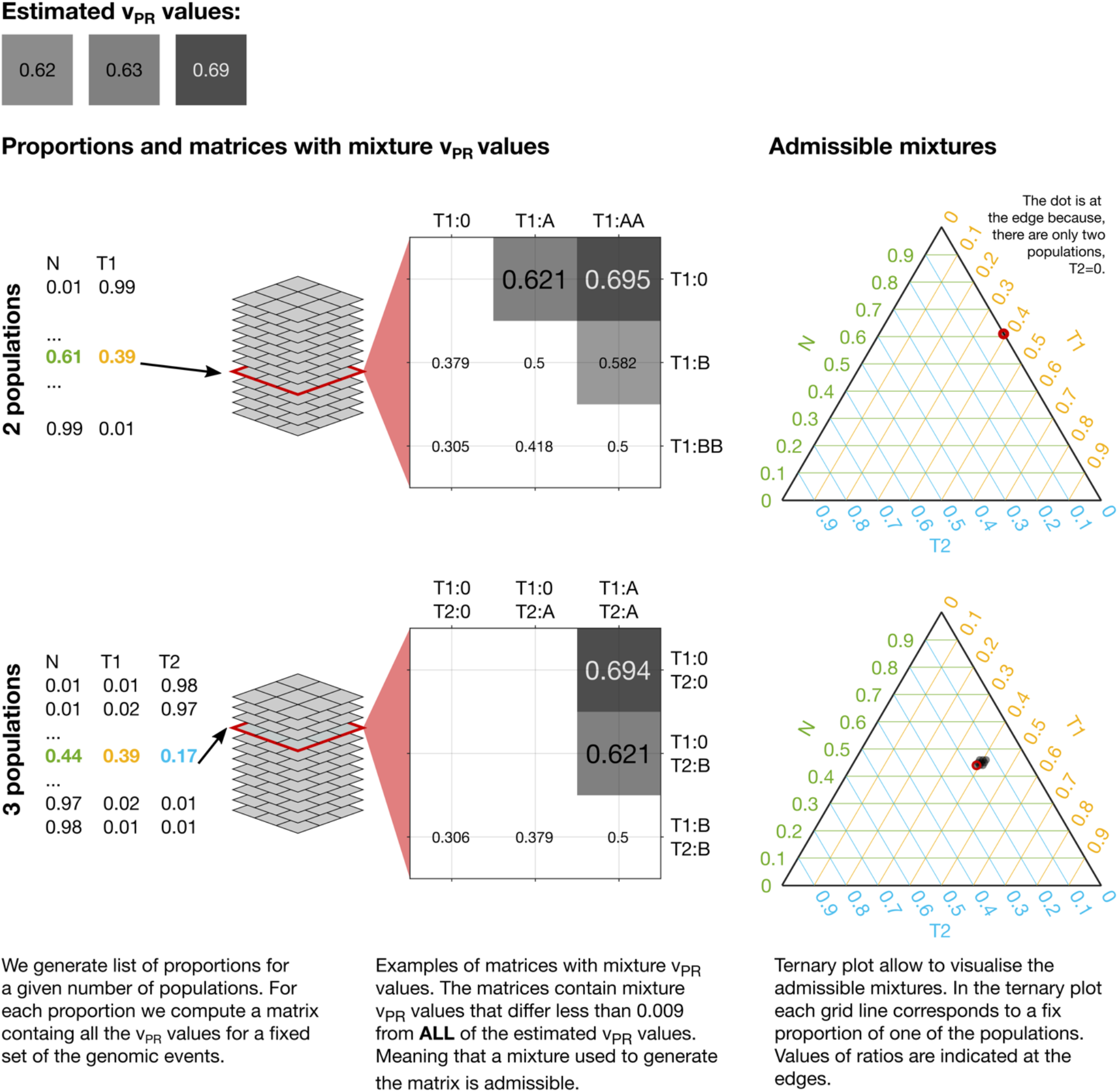
Mixtures admissible by the V_PR_ values estimated from data. To uncover mixtures that could produce the three estimated V_PR_ values we perform an exhaustive approximate search of all the possible V_PR_ values produced by any mixture of the populations with a given set of genetic events. In each case we generate a full set of proportions with a given step (e.g. 0.01) and compute all the possible V_PR_ values that such a mixture could produce. In the illustrated cases: 2 populations (1 tumour) could produce the estimated V_PR_ values through a deletion (estimated V_PR_=0.62 and V_PR_=0.63) and via deletion of one allele and duplication of another (estimated V_PR_=0.69); 3 populations (2 tumours) could produce the estimated V_PR_ values through a deletion in one of the tumour populations (estimated V_PR_=0.62 and V_PR_=0.63) and via deletion in both of the tumour populations (estimated V_PR_=0.69). The 2 populations case admits a single mixture and the 3 populations allow 9 mixtures with similar compositions. The admissible mixtures are depicted on the ternary plots, red circle indicates solution corresponding to the presented matrix. We exclude mixture V_PR_ values that result from deletion of both the variant and reference alleles (empty fields in the matrices).

To represent the admissible mixtures graphically we use ternary plots. Ternary plot allows to illustrate composition of three components using just two dimensions. The composition, represented by ratios of the three components which sum to a constant, is depicted as point inside or on the edge of an equilateral triangle. If the point is on the edges, the composition has only two components. To facilitate interpretation of the ternary plots, we also plot the grid lines that are parallel to the sides of the triangle. These gridlines indicate the directions of constant ratios of the components. Along such direction the ratio of one of the components is fixed and only the other two ratios vary. Examples of visualisation of admissible mixtures on ternary plots are shown in Figs. 5 and 6.

**Fig. 6.**
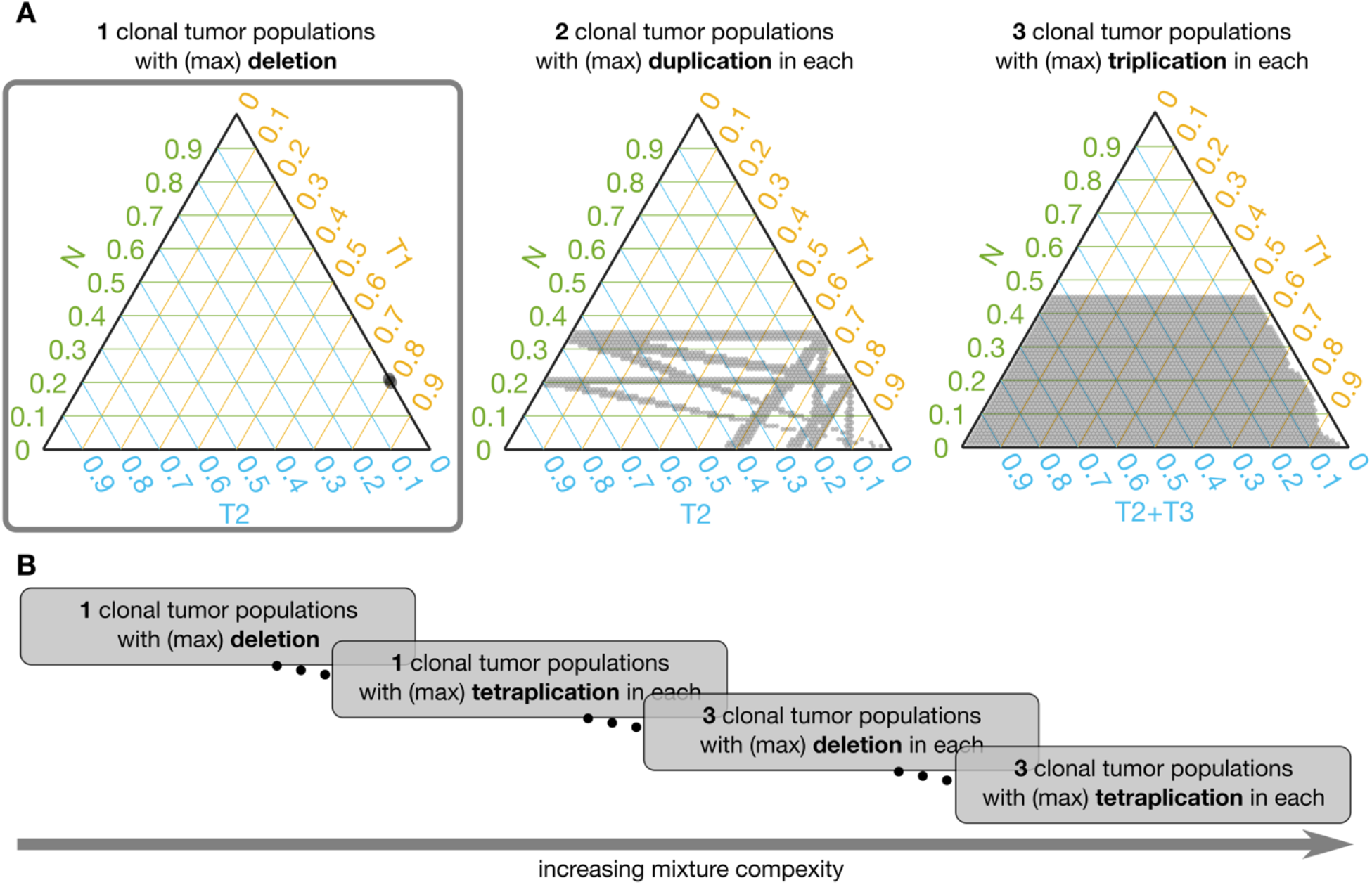
Admissible mixtures for increasing mixture complexity. **A** shows admissible mixtures for 3 different mixtures with increasing complexity. The simplest mixture (mixture with the lowest number of components and the simplest set of genetic events) is show within the grey frame. On each ternary plot, the admissible mixtures are indicated by grey dots. The green axis indicates proportion of the normal population (N), the yellow axis indicates proportion of the 1^st^ tumour population (T1), the blue axis shows proportion of the 2^nd^ tumour population (T2) or sum of the 2^nd^ and 3^rd^ tumour populations (T2+T3). **B** Schematic representation of increasing complexity of the mixture models. From a mixture of 1 normal and 1 tumour population in which only deletion is possible to a model with 1 normal and 3 tumour populations and each can have deletions, du-, tri- and tetraplications.

To facilitate analysis of the admissible mixtures returned by the search procedure we introduce mixture complexity. Mixture complexity is a measure that increases with number of populations as well as with variety of genetic events. From the simplest mixture of 1 normal and 1 tumour population in which only deletions are possible to a model with 1 normal and multiple tumour populations where each can have deletions, and any level of multiplications. In practice, we set the limit at 3 tumour populations and tetraplications. Mixture complexity helps to group and visualise admissible mixtures. Mixtures with higher complexity allow more possible V_PR_ values, meaning that it is easier to find the match with the estimated V_PR_ values and the number of admissible mixtures increases (see Figure 6). We, further, observe that proportion of normal population, p_N_, increases with a number of clonal tumour populations included in the model mixture and that, generally, p_N_ stays constant with increasing variety of genetic events, for a fixed number of clonal tumour populations.

### 2.2 Analysis of RNA-DNA relationships

GetAllele is readily applicable to assess RNA-DNA relationships between normal and tumour sequencing signals derived from the same sample/individual (matched datasets). As a proof of concept, we assessed matched normal and tumour exome and transcriptome sequencing data of 72 breast carcinoma (BRCA) datasets with pre-assessed copy-number and genome admixture estimation acquired through TCGA (Supplementary Table 1). For these datasets, purity and genome admixture has been assessed using at least three of the following five approaches: ESTIMATE, ABSOLUTE, LUMP, IHC, and the Consensus Purity Estimation (CPE)(Aran, et al., 2015; Carter, et al., 2012; Katkovnik, et al., 2002; Pagès, et al., 2010; Yoshihara, et al., 2013; Zheng, et al., 2014). In addition, on the same datasets we applied THetA – a popular tool for assessing CNA and admixture from sequencing data (Oesper, et al., 2013; Oesper, et al., 2014).

#### 2.2.1 Segmentation results

Segmentation of the data, based on the tumour exome signal, resulted in 2697 windows across the 72 datasets. We excluded from further analysis 294 windows where either tumour exome or transcriptome had V_PR_>=0.58 but their VAF distribution could not be differentiated from the model VAF distributions with V_PR_=0.5 (p>1e-5, Kolmogorov Smirnov test, equivalent to Bonferroni FWER correction for 100000 comparisons). The 294 excluded windows correspond to 4% of the data in terms of number of base pairs in the windows and 4% of all the available data points; i.e. they are short and contain only few VAF values. In the remaining 2403 windows, we systematically examined the similarity between corresponding VAF_TEX_ (tumour exome), VAF_TTR_ (tumour transcriptome) and CNA. We obtained several distinct patterns of coordinated RNA-DNA allelic behaviour as well as correlations with CNA data.

In 60% of all analysed windows the distributions of VAF_TEX_ and VAF_TTR_ were statistically concordant (had p>1e-5, Kolmogorov Smirnov test), and in 40% they were statistically discordant (p<1e-5, Kolmogorov Smirnov test). In 2 windows VAF_TEX_ and VAF_TTR_ had the same V_PR_ but had statistically different distributions (p<1e-5, Kolmogorov Smirnov test), we consider such windows as concordant; Kolmogorov-Smirnov test is very sensitive for differences between distributions, V_PR_ fitting is more robust. In the vast majority of the discordant windows V_PR_ of the VAF_TTR_, V_PR,TTR_, was higher than V_PR_ of the VAF_TEX_, V_PR,TEX_, (only in 21 out of 959 discordant windows V_PR_,TTR was lower than V_PR,TEX_).

#### 2.2.2 Concurrence of segmentation based on DNA and RNA

We next analysed the concurrence between windows resulting from independent segmentations of the dataset based on the tumour exome and transcriptome signals in the datasets (2697 and 3605 windows, respectively, across all the samples). We first assessed chromosome-wise alignment of the start and end points of the windows. In 45% of the chromosomes both VAF_TEX_ and VAF_TTR_ signals produce a single window that contains the whole chromosome. In 33% of chromosomes both signals produced multiple windows. These windows are well aligned, with 90% of the break-points within 7% difference in terms of the number of data points in the chromosome (Q50=0.02%, Q75=2% of data points in the chromosome). The probability of observing such an alignment by chance is smaller than p=1e-5 (100,000 bootstrap samples with breaking points assigned randomly in all the individual chromosomes where both signals produced multiple windows). In 22% of the chromosome windows based on VAF_TEX_ and VAF_TTR_ the signals were positionally discordant – one signal produced a single window containing whole chromosome while the other produced multiple windows.

To compare the V_PR_ values in the 55% of chromosomes where at least one signal produced more than one window, we computed chromosome-wise mean absolute error (MAE) between the V_PR_ in two sets of windows. To account for different start and end points of the windows we interpolated the V_PR_ values (nearest neighbour interpolation) at each data point in the chromosome. We separately compared the V_PR,TEX_ and V_PR,TTR_ values. The alignment in terms of MAE is very good, V_PR,TEX_ agreed perfectly in 11% of the chromosomes and had the percentiles of MAE equal to Q50= 0.012, Q75= 0.022 and Q97.5= 0.047, while V_PR,TTR_ agreed perfectly in 8% but had slightly higher percentiles of MAE Q50= 0.019, Q75= 0.034 and Q97.5= 0.07. V_PR,TEX_ and V_PR,TTR_ values had MAE=0 simultaneously in 4% of the chromosomes. Probability of observing such values of MAE by chance is smaller than p=1e-3 (1000 random assignments of V_PR,TEX_ and V_PR,TTR_ values to windows in the 873 chromosomes where at least one signal had more than one window). It is noteworthy that MAE Q97.5<0.07 is comparable with the confidence interval of single V_PR_ estimate based on 50 VAF values. In other words, both signals in a sample (Tex and Ttr) give very similar results in terms of windows’ segmentation and estimated values of the V_PR_. Albeit, segmentation of VAF_TTR_ generates a higher number of windows. The higher number of VAF_TTR_ windows indicates that transcriptional regulation occurs at a smaller scale than allelic status changes in DNA. Figure 7 shows examples of concurrence between windows based on VAF_TEX_ and VAF_TTR_ signals in a positionally concordant chromosome (both signals produced multiple windows).

**Fig. 7.**
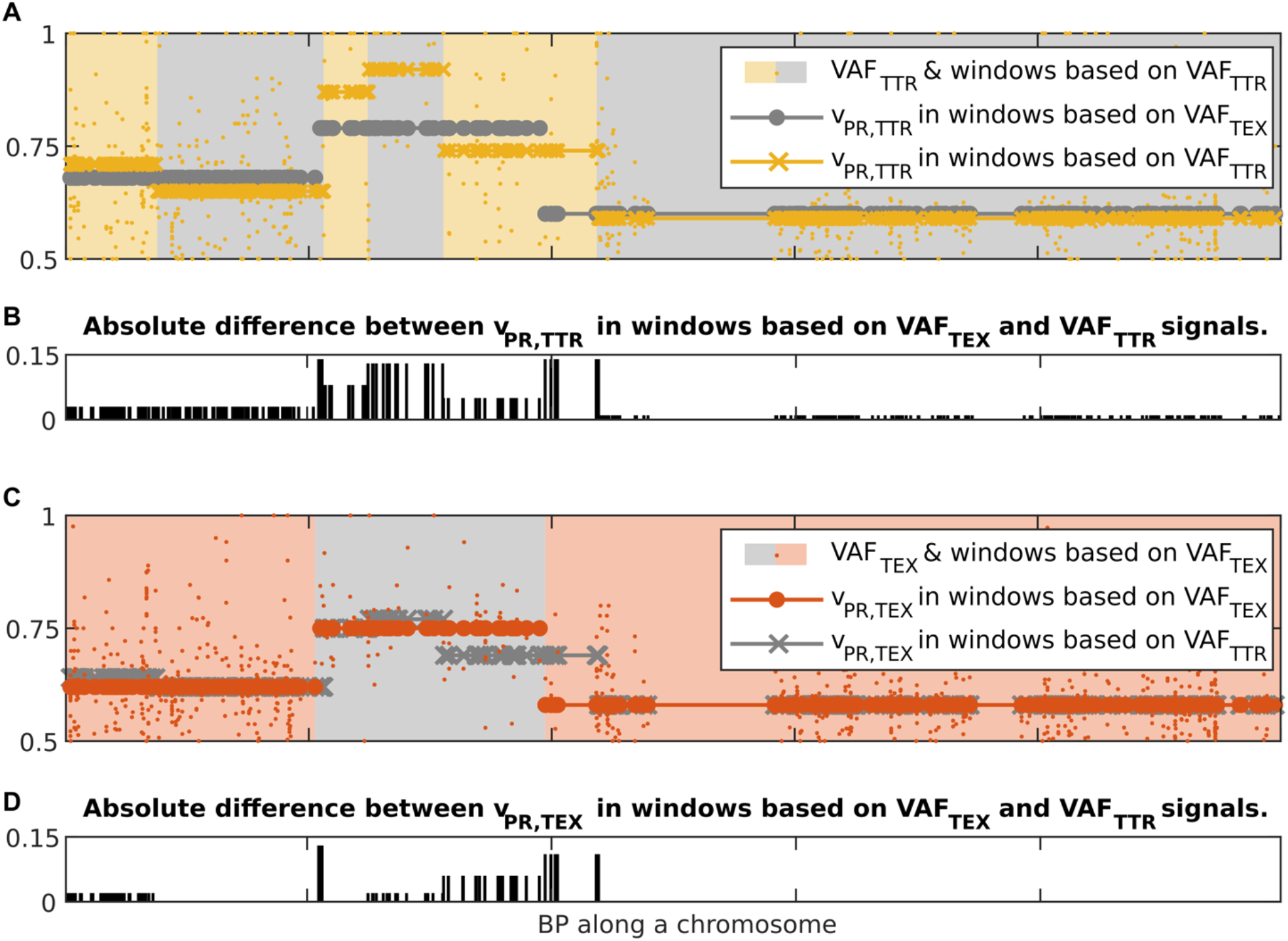
Illustration of concurrence between windows resulting from independent segmentations of the dataset based on the VAF_TEX_ and VAF_TTR_ signals. **A** yellow dots, VAF_TTR_; grey circles, V_PR,TTR_ interpolated at all data points in windows based on VAF_TEX_; yellow crosses, V_PR,TTR_ interpolated at all data points in windows based on VAF_TTR_. **B** bar plot of the absolute difference between the V_PR_ values in the two kinds of windows. **C** orange dots, VAF_TEX_; grey crosses, V_PR,TEX_ interpolated at all data points in windows based on VAF_TEX_; orange dots V_PR,TEX_ interpolated at all data points in windows based on VAF_TTR_. **D** bar plot of the absolute difference between the V_PR_ values in the two kinds of windows.

#### 2.2.3 Correlation between V_PR_ and CNA

Finally, we assess the correlations between V_PR_ and CNA in the individual datasets. We separately computed correlations for deletions and amplifications. In order to separate deletions and amplifications, for each data set we found CNA_MIN_, value of the CNA in the range −0.3 to 0.3 that had the smallest corresponding V_PR,TEX_. To account for observed variability of the CNA values near the CNA_MIN_, we set the threshold for amplifications to CNA_A_ = CNA_MIN_-0.05, and for deletions we set it to CNA_D_ = CNA_MIN_+0.05.

For VAF_TEX_ we observed significant correlations with negative trend between V_PR,TEX_ and CNA ≤ CNA_D_ in 57 datasets and with positive trend between V_PR_,TEX and CNA ≤ CNA_A_ in 39 datasets (pFDR<0.05, Pearson’s correlation with Benjamini Hochberg multiple comparison correction for 72 samples). For VAF_TTR_ we observed significant correlations with negative trend between V_PR,TTR_ and CNA ≤ CNA_D_ in 62 datasets and with positive trend between V_PR,TTR_ and CNA ≥ CNA_A_ in 33 datasets (pFDR<0.05, Pearson correlation with Benjamini Hochberg correction). These correlations indicate that although V_PR_,TEX as well as V_PR,TTR_ values capture information contained in CNA, they do not differentiate between positive and negative values of the CNA.

Figure 8 shows four typical patterns of correlation between the CNA and V_PR_ values observed in the data. In Figure 8A all the values of CNA are close to CNA_MIN_. In Figure 8B the relationship between CNA and V_PR_ is noisy, only correlation between V_PR,TTR_ and CNA ≤ CNA_D_ are statistically significant (r_TEX,CNA,D_ = −0.29, p_FDR_ = 0.063; r_TEX,CNA,D_ = −0.38, p_FDR_ = 0.012; r_TEX,CNA,A_ = 0.14, p_FDR_ = 0.58; r_TEX,CNA,A_ = 0.19, p_FDR_ = 0.47; Pearson’s correlation with Benjamini Hochberg multiple comparison correction for 72 samples). In Figure 8C all the correlations are statistically significant, V_PR_,TTR values (circles) follow closely the V_PR_,TEX (squares) indicating that in most of the windows distributions of the VAF_TEX_ and VAF_TTR_ are concordant (r_TEX,CNA,D_ = −0.91, p_FDR_<1e-10; r_TEX,CNA,D_ = −0.96, p_FDR_<1e-10; r_TEX,CNA,A_ = 0.92, p_FDR_<1e-10; r_TEX,CNA,A_ = 0.95, p_FDR_<1e-10). In Figure 8D correlations between V_PR_,TEX, V_PR_,TTR and CNA ≤ CNA_D_ are statistically significant, but there is a big difference (with median of 0.18) between V_PR_,TEX and V_PR_,TTR values, indicating that in most of the windows the distributions of the VAF_TEX_ and VAF_TTR_ in this dataset are discordant (r_TEX,CNA,D_ = −0.44, p_FDR_ = 0.047; r_TEX,CNA,D_ = −0.64, p_FDR_ = 0.0017; r_TEX,CNA,A_ = 0.44 p_FDR_ = 0.16; r_TEX,CNA,A_ = 0.28, p_FDR_ = 0.41). In many of the datasets we observe that the V_PR_,TTR values are higher than the corresponding V_PR,TEX_ values (median V_PR,TTR_–V_PR,TEX_ = 0.03). Correlations between V_PR_ and CNA in all datasets are shown in the Supplementary Figure 1.

**Fig. 8.**
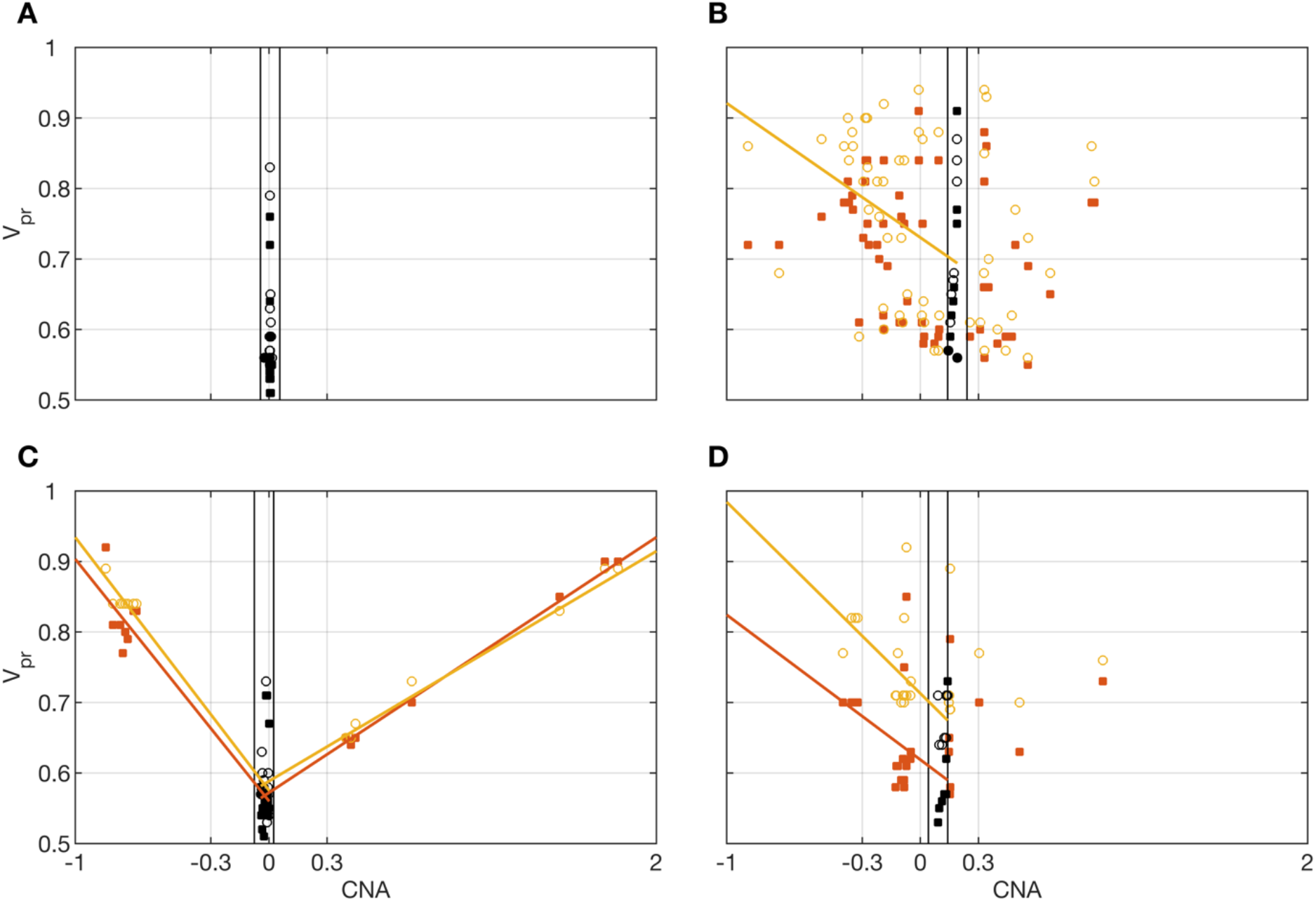
Illustration of the correlations between V_PR_ and CNA. Orange squares V_PR,TEX_, yellow circles V_PR,TTR_. Lines, least-squares fitted trends for significant correlations (orange correlation with V_PR,TEX_, yellow correlation with V_PR,TTR_). Black, V_PR_ for CNA_MIN_±0.05. Correlations for all the datasets are shown in Supplementary Figure 1. **A** all the values of CNA are close to CNA_MIN_=0. **B** relationship between CNA and V_PR_ is noisy, only some correlations are statistically significant. **C** all the correlations are statistically significant, V_PR,TTR_ values (circles) follow closely the V_PR,TEX_ (squares) indicating concordance of the VAF_TEX_ and VAF_TTR_ distributions. **D** only correlations for CNA ≤ CNA_D_ are statistically significant.

### 2.3 V_PR_ based purity estimation

To demonstrate a practical application of estimating the admissible mixtures by means of comparing the estimated V_PR,TEX_ values with the mixture V_PR_ values we next use them to estimate purity of the samples. To this end we compared the V_PR_ based purity (VBP) estimates with ESTIMATE, ABSOLUTE, LUMP, IHC, and the Consensus Purity Estimation (CPE)(Aran, et al., 2015; Carter, et al., 2012; Katkovnik, et al., 2002; Pagès, et al., 2010; Yoshihara, et al., 2013; Zheng, et al., 2014).

To obtain the VBP estimate we used V_PR,TEX_ values. We, first, selected the V_PR,TEX_ values that: 1. are estimated with high confidence, i.e. are based on at least 50 data points; 2. are most likely heterozygous in normal exome, i.e., have a corresponding V_PR_ value in normal exome V_PR,NEX_ <0.59; 3. most likely have V_PR,TEX_ > 0.5, i.e., their p-value for comparison with V_PR,TEX_ = 0.5 is very small p<1e-5 (Kolmogorov-Smirnov test). Next, we used the selected V_PR,TEX_ values to find all admissible mixtures (with 1 to 3 tumour populations and allowing for all events, from deletions to tetraplications). To estimate the VBP, out of all the admissible mixtures we chose these with lowest mixture complexity and among these mixtures we take one with the highest pN (proportion of the normal population). The VBP, percentage of tumour populations in the sample, is then given as 1-pN. Such approach provides rather conservative estimates of VBP (the smallest 1-pN). However, GetAllele can be extended to offer alternative methods of employing the admissible mixtures to estimate VBP. Development, analysis and comparison of alternative VBP estimation methods is beyond scope of the current paper.

Figure 9A shows violin plots of all considered 1-p_N_ values and (x) indicates the smallest value taken as a VBP estimate. In two of the datasets we could not estimate the purity due to lack of suitable V_PR_,TEX values. The VBP estimates shows the best agreement with ABSOLUTE method (r=0.76, p<3.4e-14, Pearson’s correlation, Fig. 9B2). We suppose that this is because the ABSOLUTE method is based on copy number distributions, and our analysis (Section 2.2.3) revealed high correlations between the CNAs and V_PR_ values. Similarly to the ABSOLUTE method, VBP estimates are generally lower than the other purity estimates (ESTIMATE, LUMP, IHC, CPE); see Fig. 9B1-B5.

**Fig. 9.**
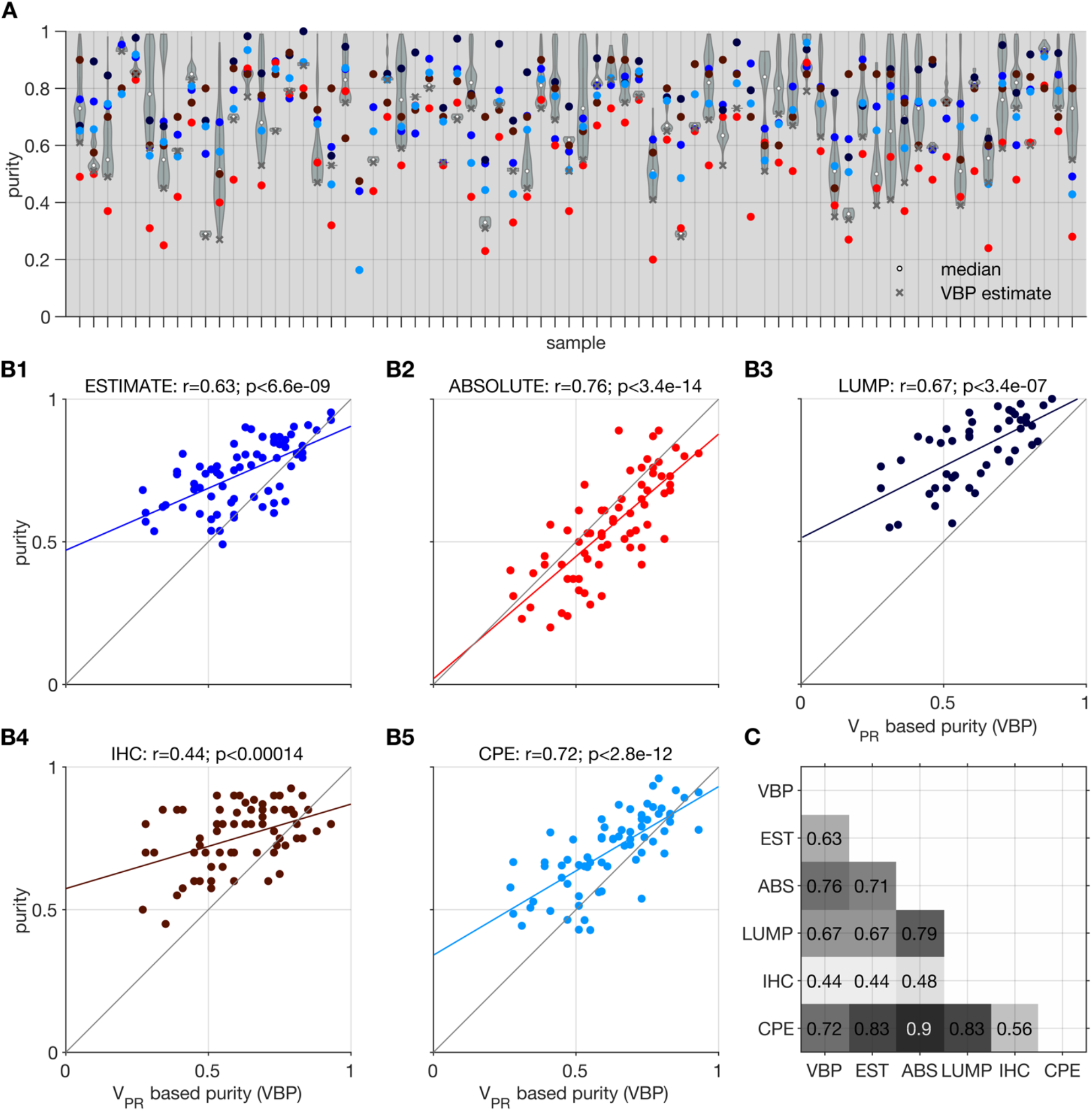
Illustration of purity estimation based on model mixtures and V_PR,TEX_. Comparison of the Estimate (EST), Absolute (ABS), LUMP, IHC, and the consensus purity estimate (CPE) methods with V_PR_ based purity (VBP). **A** violin plots show distributions of purity based on all the admissible proportions of the normal population (x) indicates the lowest value selected as the most conservative estimate; colours corresponding to the different methods are indicated in B1-5. **B1-5** Correlation of the VBP with individual methods; coloured line indicates best fit linear trend. **C.** Matrix showing significant p<0.05 Pearson’s correlation coefficients between all tested methods.

The approach presented in this section differs from other methods for inferring genomic mixture composition (e.g. SciClone (Miller, et al., 2014), PyClone (Roth, et al., 2014) or TPES (Locallo, et al., 2019)) in that it is based on changes of the alleles multiplicity estimated along continuous multi-SNV genomic regions. The V_PR_ values used to estimate admissible mixtures are based on windows with at least 50 VAF values which extend over millions of base pairs. In this way, the presented method is complementary to the SciClone and PyClone methods that use VAFs of somatic mutations.

## 3 Discussion

Potential for integrative analysis of RNA and DNA sequence data is facilitated by the growing availability of RNA and DNA sequencing datasets and by the technological advances now enabling simultaneous RNA and DNA sequencing from the same source (Macaulay, et al., 2016; Reuter, et al., 2016; The, et al., 2012). However, integrative RNA and DNA analyses are challenged by limited compatibility between RNA and DNA datasets and high technical variance of the sequencing-produced signals. Our approach – GeTallele – addresses the compatibility by restricting the analyses to confidently co-covered DNA and RNA regions, and the high variability - through quantification of differences between the DNA and RNA signals.

In contrast to other methods based on statistical modelling, GeTallele is based on a mechanistic model of VAF distributions that depends on the distribution of total reads (extracted from data) and the V_PR_ parameter. The brute-force simplicity and transparency of the presented methodology is one of its biggest advantages. Additionally, in contrast to other methods, to analyse sequencing data GeTallele uses data segments that include multiple adjacent SNVs. The proposed mechanistic model indicates that due to probabilistic nature of the reads that estimates of genomic events based on continuous multi-SNV regions are intrinsically more robust than estimates based on a single SNV. Finally, GeTallele offers a unified pipeline to evaluate and visualise DNA-RNA whereas many of the other methods cited herein are used in conjunction with each other to achieve a similar result.

Using GeTallele, we detected several relationships between DNA-RNA allele frequencies and biological processes. First, in chromosomes affected by deletions and amplifications, VAF_RNA_ and VAF_DNA_ showed highly concordant breakpoint calls. This indicates that VAF_RNA_ alone can serve as preliminary indicator for deletions and amplifications, which can facilitate the applications of RNA-sequencing analysis on the large and constantly growing collections of transcriptome sequencing data. Second, we showcased that V_PR,TEX_ can be used to estimate sample composition (in terms of proportions of normal and tumour populations) and as a result its purity. Once the mixture composition is decided by the user, GeTallele allow to further interrogate genetic events in each population at a specific data segment.

## 4 Conclusions

Based on our results, variant probability V_PR_ can serve as a dependable indicator to assess gene and chromosomal allele asymmetries and to aid calls of genomic events in matched sequencing RNA and DNA datasets. Methods for estimating and analysing V_PR_ values are implemented in a GeTallele toolbox. GeTallele provides a singular suit of functions for integrative analysis, statistical assessment and visualization of the observed patterns at desired resolution, including chromosome, gene, or custom genome region.

## 5 Methods

### 5.1 Samples

The GeTallele was developed using sequencing datasets from paired normal and tumour tissue obtained from 72 female patients with breast invasive carcinoma (BRCA) from The Cancer Genome Atlas (TCGA). Each of the 72 datasets contains four matched sequencing datasets: normal exome (Nex), normal transcriptome (Ntr), tumour exome (Tex), and tumour transcriptome (Ttr). In addition, we required each tumour sample to have at least three of the following five purity estimates - Estimate, Absolute, LUMP, IHC, and the consensus purity estimate (CPE), (Supplementary Table 1). Finally, each sample was required to have CNA estimation (genomic segment means based on Genome-Wide-SNPv6 hybridization array) (Aran, et al., 2015; Carter, et al., 2012; Katkovnik, et al., 2002; Pagès, et al., 2010; Yoshihara, et al., 2013; Zheng, et al., 2014).

### 5.2 Data processing

All datasets were generated through paired-end sequencing on an Illumina HiSeq platform. The human genome reference (hg38)-aligned sequencing reads (Binary Alignment Maps, .bams) were downloaded from the Genomic Data Commons Data Portal (https://portal.gdc.cancer.gov/) and processed downstream through an in-house pipeline. After variant call (Li, 2011), the RNA and DNA alignments, together with the variant lists were processed through the read count module of the package RNA2DNAlign (Movassagh, et al., 2016), to produce variant and reference sequencing read counts for all the variant positions in all four sequencing signals (normal exome, normal transcriptome, tumour exome and tumour transcriptome). Selected read count assessments were visually examined using Integrative Genomics Viewer (Thorvaldsdóttir, et al., 2013).

### 5.3 Data segmentation

To analyse variant allele frequencies (VAF) at genome-wide level, GeTallele first divides the VAF dataset into a set of non-overlapping windows along the chromosomes. Segmentation of the dataset into windows is based on a sequencing signal chosen out of all the available datasets in the aggregated aligned VAF dataset (one out of four in the presented analysis).

To partition the data into the windows GeTallele uses a parametric global method, which detects the breakpoints in the signals using its mean, as implemented in the Matlab function `findchangepts` (Killick, et al., 2012; Lavielle, 2005) or in R (Killick and Eckley, 2014). In each window, the VAF values of the chosen signal have a mean that is different from the mean in the adjacent windows. Sensitivity of breakpoint detection can be controlled using parameter `MinThreshold`, in the presented analysis it was set to 0.2. For segmentation and analysis (without loss of generality) we transform all the original VAF values to VAF=|VAF-0.5|+0.5.

### 5.4 Statistics

To test statistical significance, GeTallele uses parametric and non-parametric methods and statistical tests (Corder and Foreman, 2014; Hollander, et al., 2013). Namely, to compare distributions of the variant allele frequencies (VAF) we use Kolmogorov–Smirnov test (examples of VAF distributions are depicted in Figs. 2 and 3). To study concurrence of windows, we use permutation/ bootstrap tests. To test relations between V_PR_ and copy number alterations (CNA) we use Pearson’s correlation coefficient.

To account for multiple comparisons, we set the probability for rejecting the null hypothesis at p<1e-5, which corresponds to Bonferroni (Dunn, 1961) family-wise error rate (FWER) correction against 100000 comparisons. We use a fixed value, rather than other approaches, to ensure better consistency and reproducibility of the results. Alternatively, we apply Benjamini and Hochberg (Benjamini and Hochberg, 1995) false discovery rate (FDR) correction with a probability of accepting false positive results p_FDR_<0.05. We specify the method used in the text when reporting the results.

## List of abbreviations

BRCA: breast invasive carcinoma
CDF: cumulative distribution function
CNA: copy number alterations
CNA_D_: copy number alterations corresponding to deletions (see section 2.2.3)
CNA_A_: copy number alterations corresponding to amplifications (see section 2.2.3)
CPE: consensus purity estimate
DNA: genome
EMD: earth mover’s distance
FWER: family-wise error rate
FDR: false discovery rate
MEA: mean absolute error
Nex: normal exome
Ntr: normal transcriptome
r_TEX,CNA,D_: Pearson’s correlation coefficient between V_PR_,TEX and CNA_D_
r_TEX,CNA,A_: Pearson’s correlation coefficient between V_PR_,TEX and CNA_A_
r_TTR,CNA,D_: Pearson’s correlation coefficient between V_PR_,TTR and CNA_D_
r_TTR,CNA,A_: Pearson’s correlation coefficient between V_PR_,TTR and CNA_A_
p_FDR_: p-value after multiple comparisons Benjamini and Hochberg false discovery rate correction
PDF: probability density function
QN (e.g. Q50): N-th percentile
RNA: transcriptome
SNV: single-nucleotide variant
TCGA: the cancer genome atlas
Tex: tumour exome
Ttr: tumour transcriptome
VAF: variant allele frequency
VAF_TEX_: variant allele frequency in tumour exome sequence
VAF_TTR_: variant allele frequency in tumour transcriptome sequence
VBP: V_PR_ based purity
V_PR_: variant probability
V_PR,TEX_: variant probability estimated from tumour exome sequence
V_PR,TTR_: variant probability estimated from tumour transcriptome sequence

## Declarations

### Ethics approval and consent to participate

Not applicable

### Consent for publication

Not applicable

### Availability of data and materials

The datasets used and/or analysed during the current study are available from the corresponding author on reasonable request. GeTallele is implemented as a Matlab toolbox available at: https://github.com/SlowinskiPiotr/GeTallele. We are working on R implementation of GeTallele.

### Competing interests

The authors declare that they have no competing interests.

### Funding

This work was supported by McCormick Genomic and Proteomic Center (MGPC), The George Washington University; [MGPC_PG2018 to AH]. Work of PS was generously supported by the Wellcome Trust Institutional Strategic Support Award [204909/Z/16/Z]. KTA gratefully acknowledges the financial support of the EPSRC via grant EP/N014391/1.

### Authors’ contributions

PS, ML, PR, NA, LFS, CM, KTA, AH conception and design of the work; ML data acquisition; PS data analysis; PS, ML, PR, LFS, KTA, AH interpretation of data; PS the creation of new software used in the work; PS, ML, PR, LFS, KTA, AH have drafted the work or substantively revised it; all authors approved the submitted version. All authors agreed both to be personally accountable for the author’s own contributions and to ensure that questions related to the accuracy or integrity of any part of the work, even ones in which the author was not personally involved, are appropriately investigated, resolved, and the resolution documented in the literature.

## Acknowledgements

Not applicable

---

**Supplementary Fig. 1.**
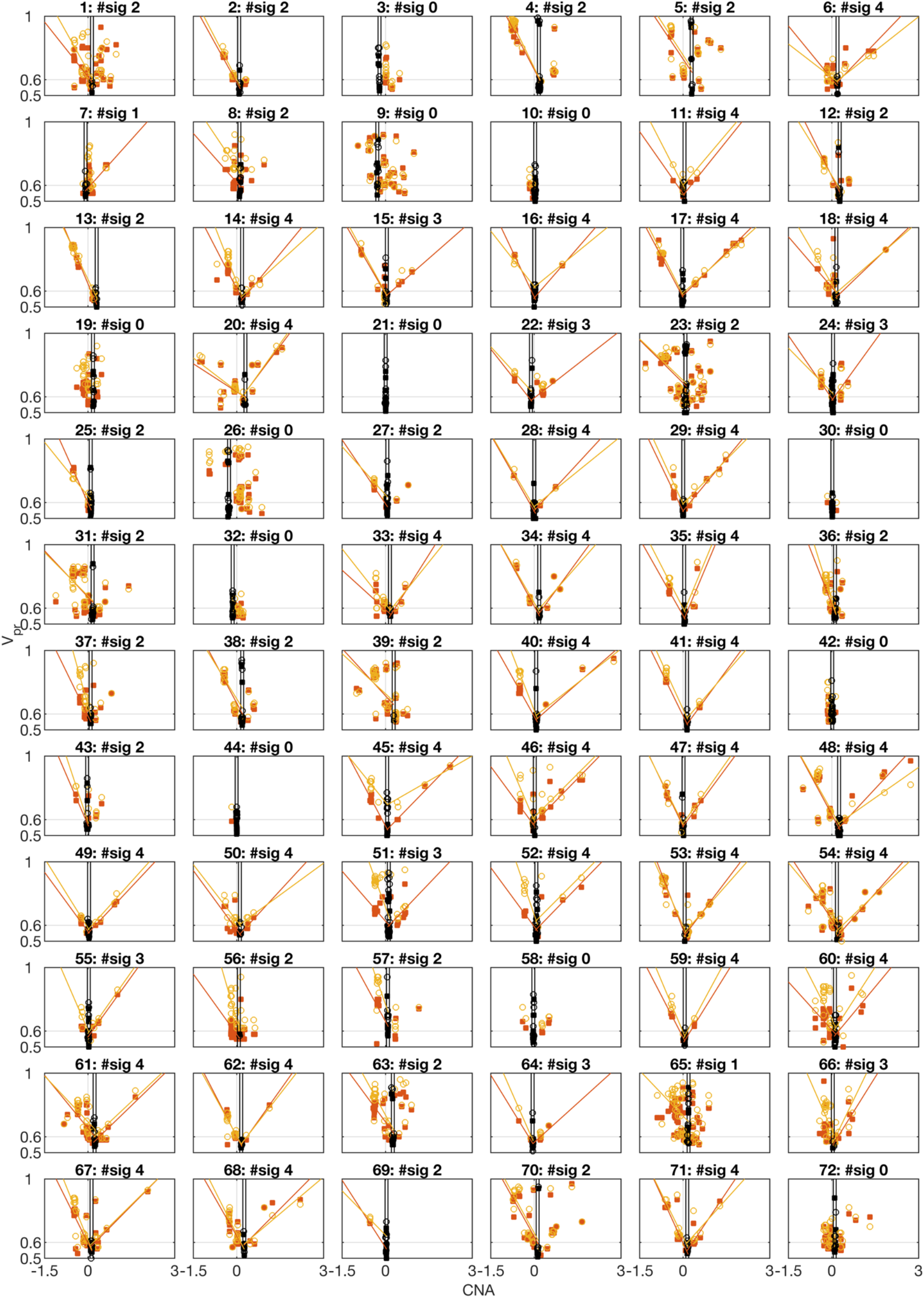
Illustration of the correlations between V_PR_ and CNA. Orange squares V_PR,TEX_, yellow circles V_PR,TTR_. Lines, leastsquares fitted trends for significant correlations (orange correlation with V_PR,TEX_, yellow correlation with V_PR,TTR_). Black, V_PR_ for CNA_MIN_±0.05. Title format Number of the dataset: #sig number of significant correlations in the dataset.

**Supplementary Table 1.**
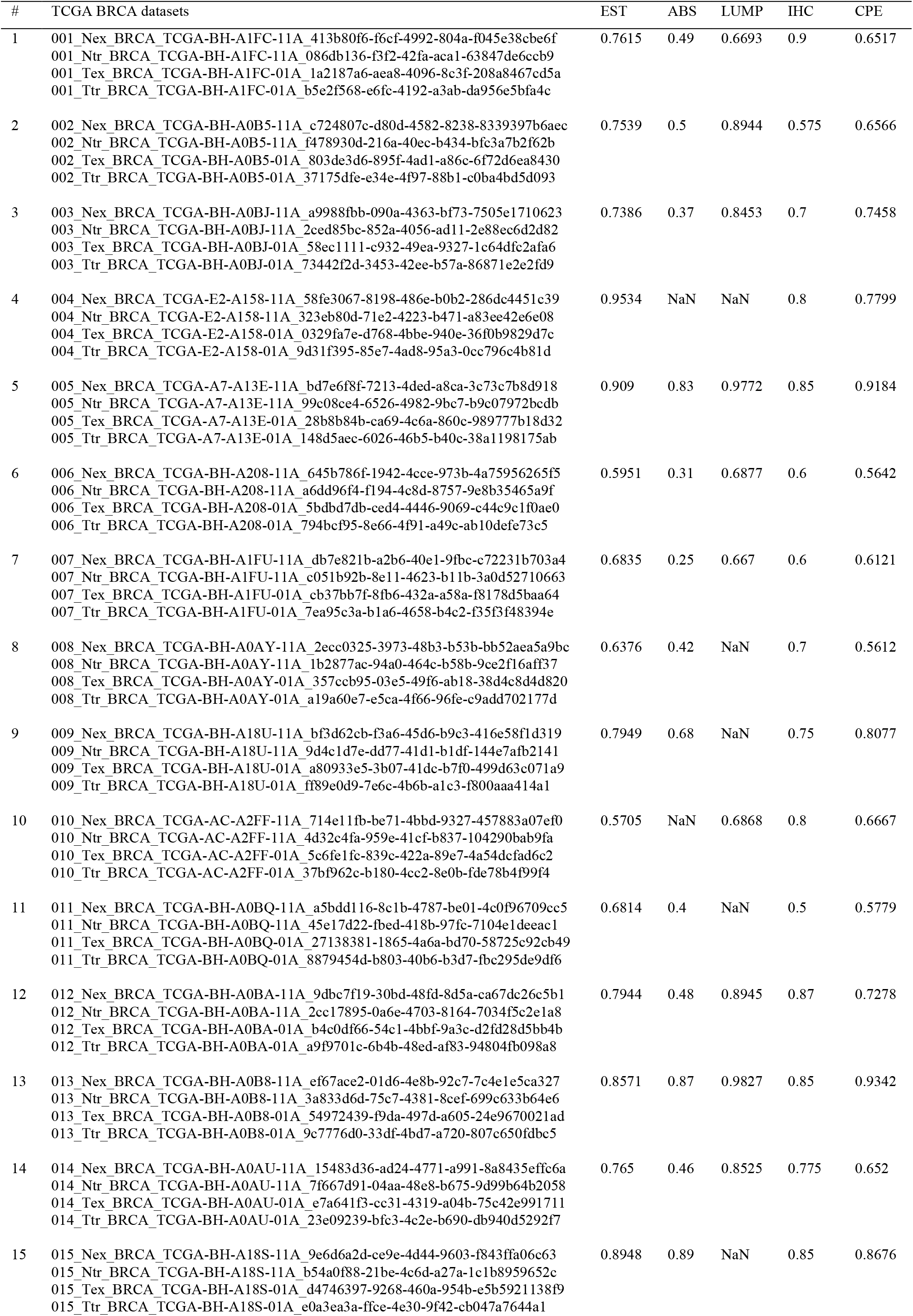
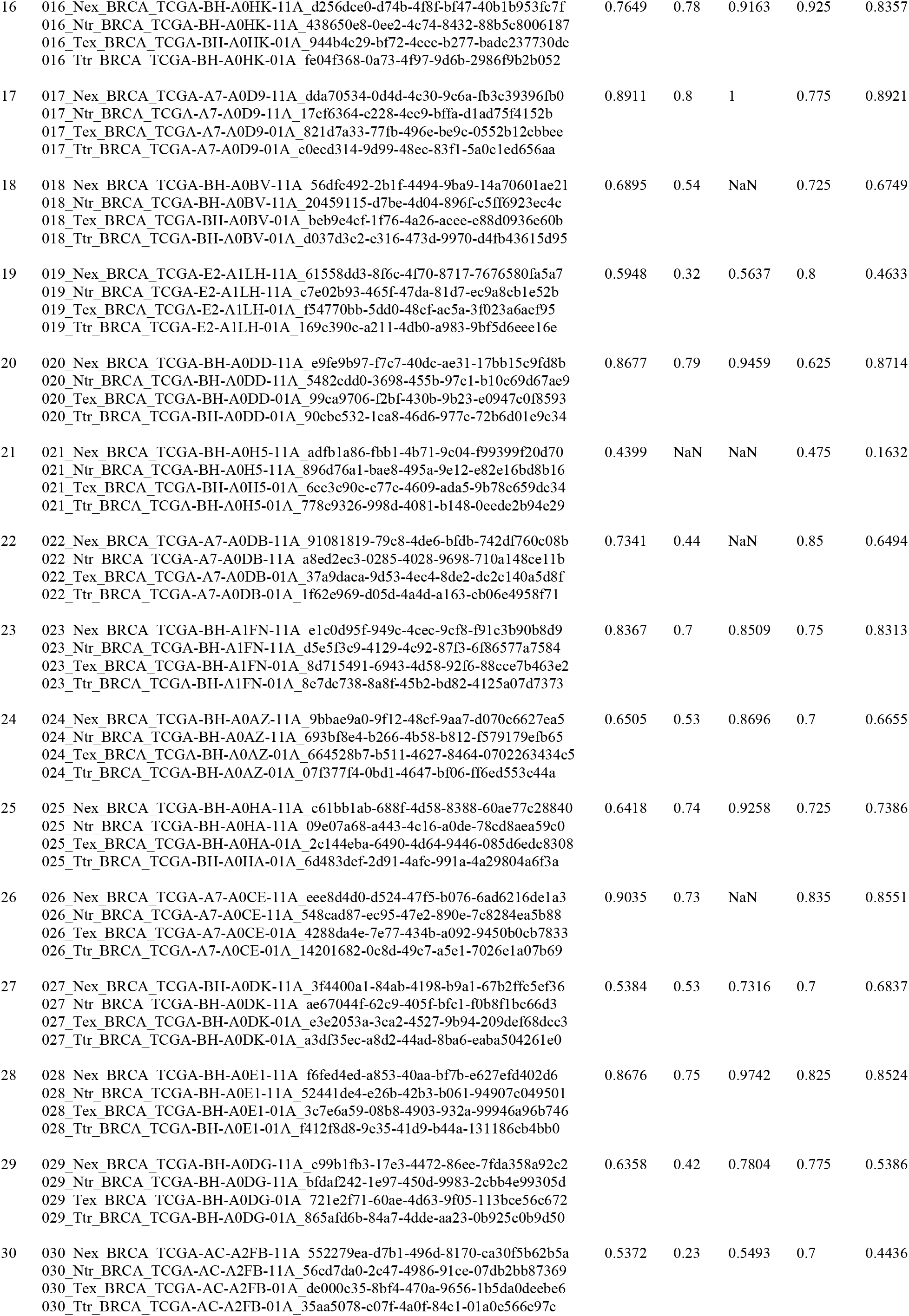
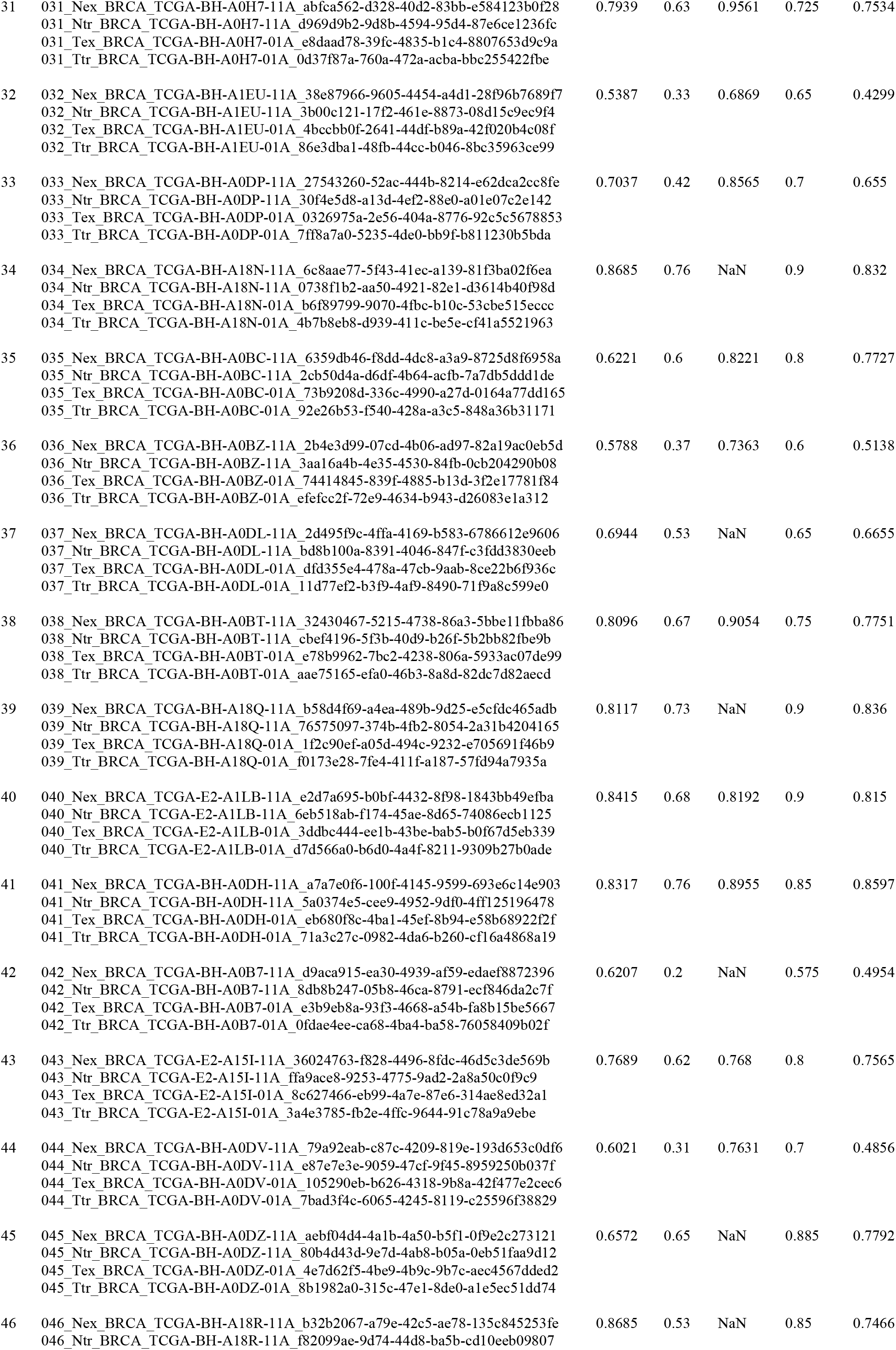
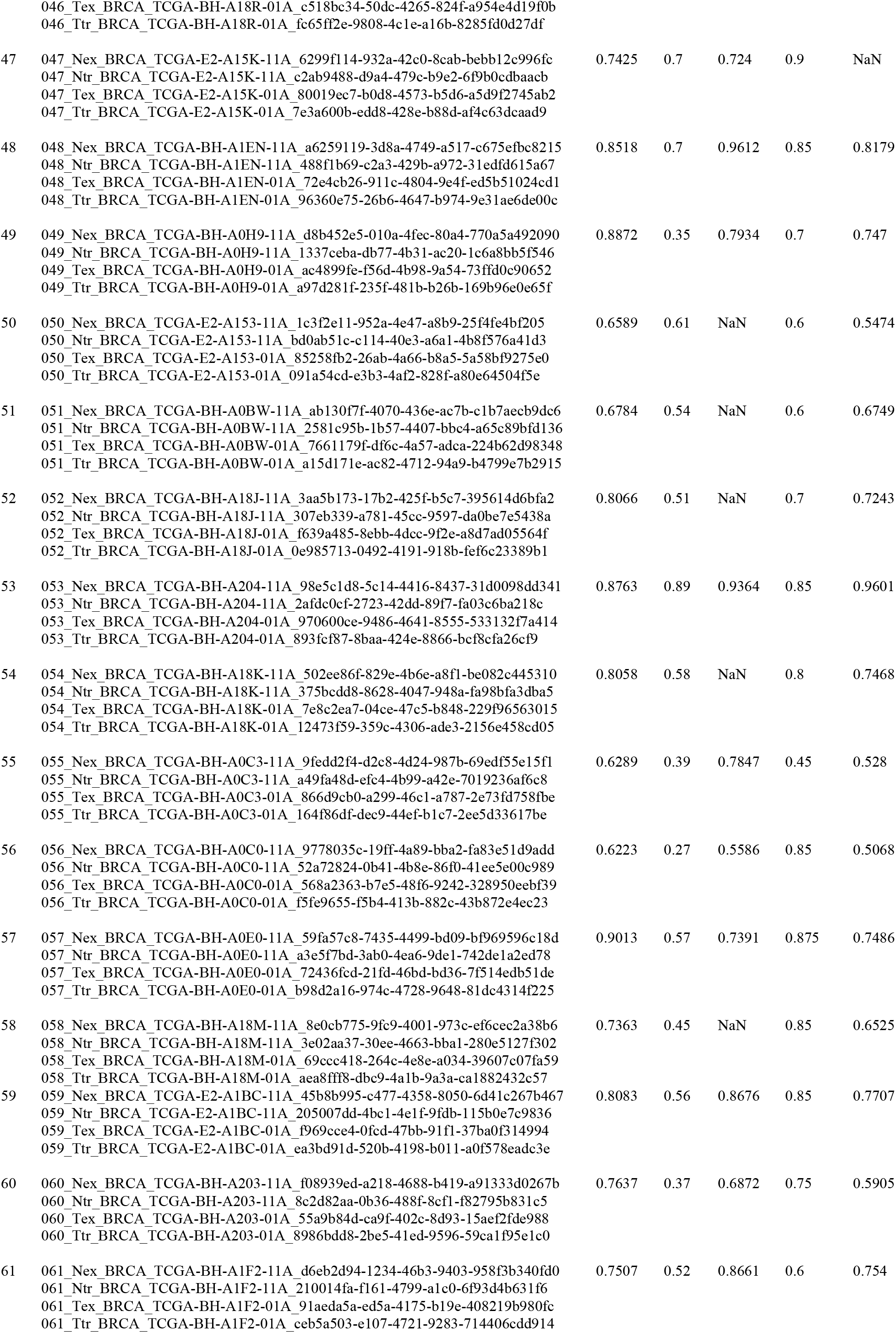
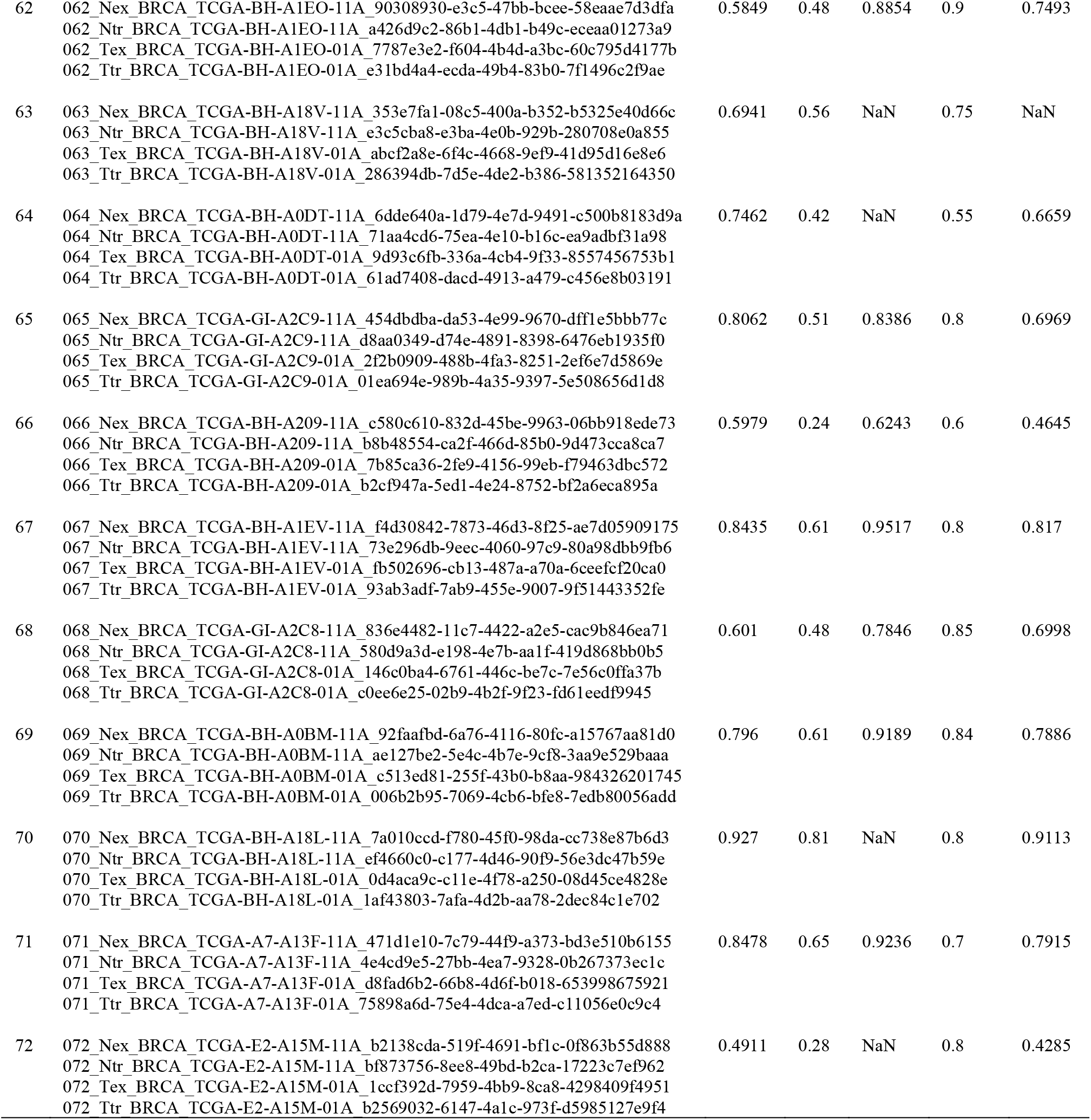
Datasets, signals and purity estimates.

## Notes

https://github.com/SlowinskiPiotr/GeTallele

### Competing Interest Statement

The authors have declared no competing interest.

